# White matter hyperintensities may be an early marker for age-related cognitive decline

**DOI:** 10.1101/2021.09.23.461560

**Authors:** Cassandra Morrison, Mahsa Dadar, Sylvia Villeneuve, D. Louis Collins, for Alzheimer’s Disease Neuroimaging Initiative

## Abstract

**Background:** Research suggests that cerebral small vessel disease (CSVD), amyloid, and pTau contribute to age-related cognitive decline. It remains unknown how these factors relate to one another, nor how they jointly contribute to cognitive decline in normal aging. This project examines the association between these pathologies and their relationship to cognitive decline in cognitively normal older adults without subjective cognitive decline.

**Methods:** A total of 230 subjects with CSF Aß42, CSF pTau181, white matter hyperintensities (WMHs) used as a proxy of CSVD and cognitive scores from the Alzheimer’s Disease Neuroimaging Initiative were included. Associations between each pathology and cognitive score were investigated using regression models. Furthermore, relationships between the three pathologies were also examined using regression models.

**Results:** At baseline, there was an inverse association between WMH load and Aß42 (*t*=-4.20, *p*<.001). There was no association between WMH load and pTau (t=0.32, p=0.75), nor with Aß42 and pTau (*t*=0.51, *p*=.61). Correcting for age, sex and education, baseline WMH load was associated with baseline ADAS-13 scores (*t*=2.59, *p*=.01) and lower follow-up executive functioning (*t*= -2.84, *p*=.005). Baseline Aß42 was associated with executive function at baseline (*t*=3.58, *p<*.004) but not at follow-up (*t*=1.05, *p*=0.30), nor with ADAS-13 at baseline (t=-0.24, p0.81) or follow-up (*t*=0.09, *p*=0.93). Finally, baseline pTau was not associated with any cognitive measure at baseline or follow-up.

**Conclusion:** Both baseline Aß42 and WMH load are associated with some baseline cognition scores, but only baseline WMH load is associated with follow-up executive functioning, indicating that it may be one of the earliest pathologies that contributes to future cognitive decline, in cognitively healthy older adults. Given that healthy older adults with WMH pathology exhibit declines in cognitive functioning, they may be less resilient to future pathology increasing their risk for cognitive impairment due to dementia than those without WMHs.

## Introduction

Alzheimer’s disease (AD) is characterized by several neuropathological brain changes such as senile plaques (ß-amyloid, Aß) and neurofibrillary tangles (Tau) deposition (Perrin et al., 2009; Sperling et al., 2011). These accumulations have been associated with the hallmarked progressive cognitive decline observed in people with AD. Both CSF Aß and tau pathologies (Donohue et al., 2017; Stomrud et al., 2010; Verberk et al., 2020) and position emission tomography (PET) Aß and tau (Aschenbrenner et al., 2018; Schöll et al., 2016; Sperling et al., 2019) have been associated with cognitive decline in healthy older adults. Tau pathology markers have been observed to be better predictors of cognitive decline than amyloid (Aschenbrenner et al., 2018; Malpas et al., 2021).

Another pathological change, cerebral small vessel disease (CSVD), results in structural damage to the white matter of the brain that is often observed as white matter hyperintensities (WMH) in T2w or FLAIR MRI in both non-clinical healthy aging populations and individuals with cognitive decline (Rhodius-Meester et al., 2017). An association between high WMH load and decreased cognitive functioning in healthy older adults is often reported. A meta-analysis observed that CSVD impacts all cognitive domains in healthy older adults, with the strongest association seen between WMH and attention and executive functioning (Kloppenborg and Geerlings, 2014). WMH load increases a healthy older adults’ risk for future development of mild cognitive impairment (MCI)(Boyle et al., 2016) and dementia (Prins et al., 2004). CSVD also continues to contribute to cognitive decline observed in AD (Kaskikallio et al., 2020) because many cases of AD have a mixed etiology (Prins and Scheltens, 2015). CSVD (as measured by WMH) has been observed to precede neurodegeneration and cognitive decline in MCI, AD, and Parkinson’s disease (Dadar et al., 2020a, 2018b), suggesting that CSVD may be involved with the etiology of cognitive decline in normal aging and cognitive impairment due to MCI and Alzheimer’s disease (AD).

While many studies have examined the effects of CSF tau and amyloid on cognitive functioning (Donohue et al., 2017; Stomrud et al., 2010; Verberk et al., 2020) and the relationship between WMH and amyloid (Dadar et al., 2020a; Hedden et al., 2012; Vemuri et al., 2015) there is limited research examining the interaction between WMH, amyloid, and tau and their joint effects on cognitive decline in healthy older adults. One study, observed no association between WMH burden and amyloid-PET but also found that increased WMH was associated with lower executive functioning whereas amyloid measured with PET had no effect on cognitive function (Marchant et al., 2012). Another study observed the association between WMHs and tau- and amyloid-PET but did not compare these pathologies to cognitive decline (Graff-Radford et al., 2019). They observed that increases in WMH were associated with increases in amyloid-PET in cognitively healthy older adults, but no relationship between tau-PET and WMH was observed (Graff-Radford et al., 2019).

One difficulty when studying these AD-related pathologies is that they occur years prior to the onset of clinical symptoms (Craig-Schapiro et al., 2009; Sperling et al., 2011), emphasizing the need to evaluate these changes in the healthy aging population. The goal of this study was thus to expand on current research by examining the relationship between cognitive changes and all three pathologies (CSVD, amyloid, and tau) in healthy older adults. The novelty of this paper was to focus on cognitively heathy older adults by excluding people with subjective cognitive decline (SCD) to avoid possible confounds caused by cognitively heathy older adults that may already be on the AD trajectory (Rabin et al., 2017). Current findings in cognitively normal elderly showing associations between the pathologies may be biased because of the inclusion of people with SCD. Another aim was to explore whether these pathologies are present in healthy older adults who are unlikely to be in the preclinical AD phase of AD. We wanted to examine how these AD-related pathologies at baseline are associated with current cognitive function and cognitive decline in healthy older adults. Furthermore, we also examined how these pathologies are related to one another.

## Methods

### Alzheimer’s Disease Neuroimaging Initiative

Data used in the preparation of this article were obtained from the Alzheimer’s Disease Neuroimaging Initiative (ADNI) database (adni.loni.usc.edu). The ADNI was launched in 2003 as a public-private partnership, led by Principal Investigator Michael W. Weiner, MD. The primary goal of ADNI has been to test whether serial magnetic resonance imaging (MRI), positron emission tomography (PET), other biological markers, and clinical and neuropsychological assessment can be combined to measure the progression of MCI and early AD. Participants were between 55 and 90 years old at the time of recruitment. The study received ethical approval from the review boards of all participating institutions. Written informed consent was obtained from participants or their study partner.

### Participants

ADNI-1 and ADNI-GO did not measure subjective cognitive decline; therefore, we applied the following study inclusion and exclusion criteria to the 406 cognitively normal controls with no SCD from the ADNI-2 and ADNI-3 cohorts with MRI scans. Full participant inclusion/exclusion criteria can be downloaded from www.adni-info.org. Briefly, Participants were between 55 and 90 years old at the time of recruitment and had no evidence of cognitive decline on either the Mini Mental Status Examination or Clinical Dementia Rating. Cognitively normal older healthy adults were considered if they were labelled as ‘Normal control’ and if they scored <16 on the Cognitive Change Index, signifying no indication of SCD. The following inclusion criteria were applied to the 406 healthy controls: having an MRI scan from which WMH load could be estimated and having both pTau and amyloid measures. Inclusion criteria also included baseline executive functioning, memory, and ADAS-13 scores. These criteria resulted in 230 participants for our study at baseline. At follow-up, 199 of these participants had cognitive scores. Figure 1 summarizes the methodology used to select participants. This project is a cohort study design that followed healthy older adults over a period of time.

**Figure 1:**
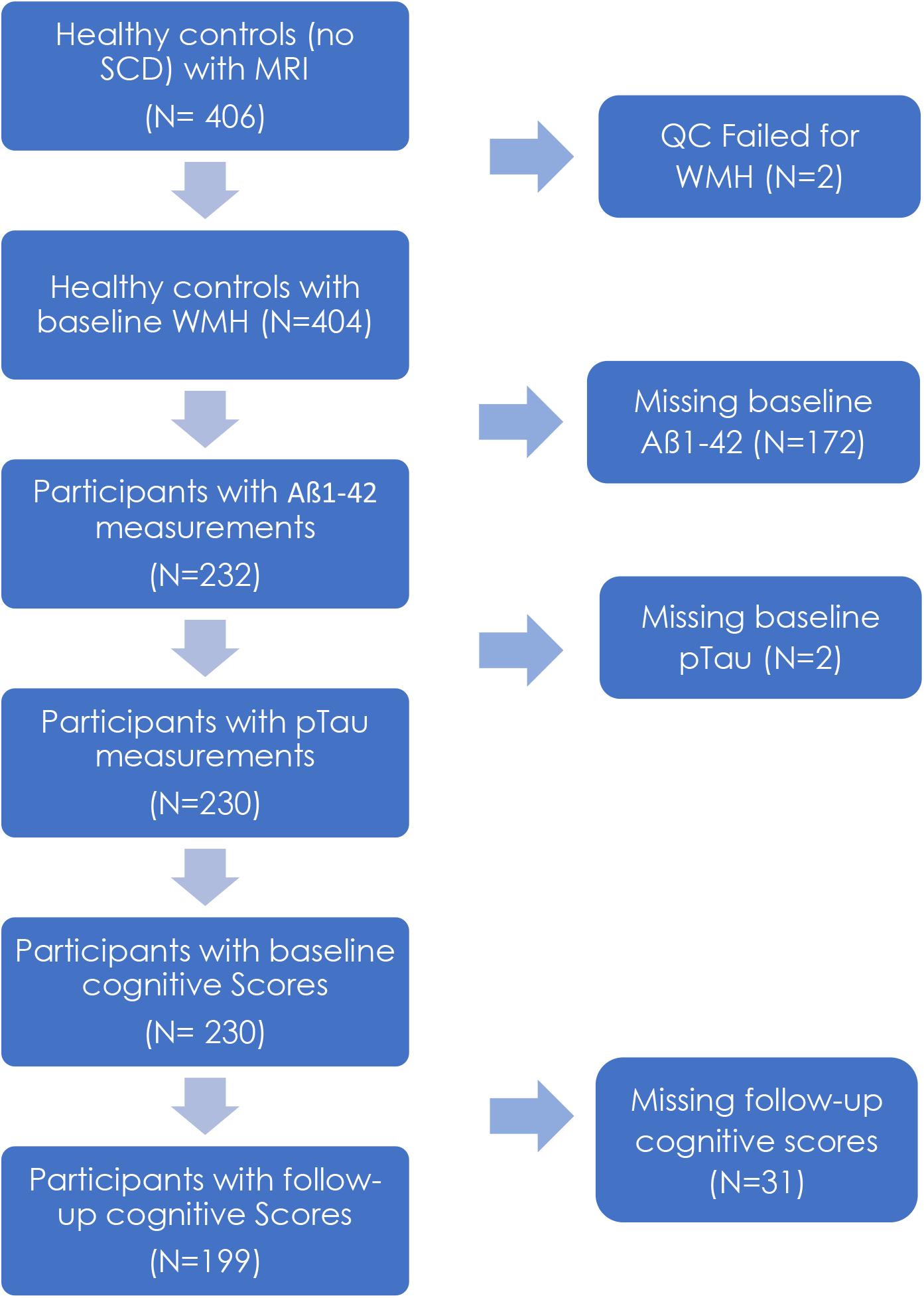
Flowchart summarizing the participant inclusion and exclusion criteria based on WMH, Aß42, pTau, and cognitive score measurements.

We used executive functioning (ADNI-EF; Gibbons et al., 2012) and memory (ADNI-MEM; Crane et al., 2012)) composite scores that have been previously developed and validated in ADNI. Alzheimer’s Disease Assessment Scale-Cog13 (ADAS-13;Mohs et al., 1997) scores were also included for all participants. The ADAS-13 measures severity of cognitive decline, for a total of 85 points, with higher scores indicating greater severity. These scores were downloaded from the ADNI public website.

To obtain cerebrospinal fluid (CSF) samples, lumbar punctions were performed as described in the ADNI procedures manual. CSF Aß42 and phosphorylated p-Tau181 (pTau) were measured using the multiplex xMAP Luminex platform (Luminex Corp, Austin, TX, USA) with the INNO-BIA AlzBio3 kit (Innogenetics) (Olsson et al., 2005; Shaw et al., 2009). The Elecsys Aß42 assay has not been established for measuring concentrations above 1700 pg/ml. Any values above this concentration were, therefore, truncated at 1700 pg/ml (Sutphen et al., 2018).

### Structural MRI acquisition and processing

All participants were imaged using a 3T scanner with T1-weighted imaging parameters (see http://adni.loni.usc.edu/methods/mri-tool/mri-analysis/ for the detailed MRI acquisition protocol). Baseline scans were downloaded from ADNI.

T1w scans for each participant were pre-processed through our standard pipeline including noise reduction (Coupe et al., 2008), intensity inhomogeneity correction (Sled et al., 1998), and intensity normalization into range [0-100]. The pre-processed images were then both linearly (9 parameters: 3 translation, 3 rotation, and 3 scaling) (Dadar et al., 2018a) and nonlinearly (1 mm^3^ grid) (Avants et al., 2008) registered to the MNI-ICBM152-2009c average template (Fonov et al., 2011). The quality of the linear and nonlinear registrations was visually verified by an experienced rater (co-author MD).

### WMH measurements

WMH measurements were obtained through a previously validated WMH segmentation technique and a library of manual segmentations based on 50 ADNI participants (independent of the 102 studied here). WMHs were automatically segmented at baseline using the T1w contrasts, along with a set of location and intensity features obtained from a library of manually segmented scans in combination with a random forest classifier to detect the WMHs in new images(Dadar et al., 2017a, 2017b). WMH load often reflects cerebrovascular disease, thus WMH load was used as a proxy for cerebrovascular pathology and was defined as the volume of all voxels identified as WMH in the standard space (in mm^3^) and are thus normalized for head size. WMH volumes were log-transformed to achieve normal distribution. This WMH technique has been developed and extensively tested specifically for assessment of WMHs in multi-center studies including validation in the ADNI cohort. For example, these techniques have been used in examining WMHs in HIV infection studies(Sanford et al., 2019) and multi-center studies including, The Parkinson’s Markers Initiative (Dadar et al., 2020b) and UK Brain bank (Dadar et al., 2020c) to assess WMHs in Parkinson’s, and the National Alzheimer’s Coordinating Center (Anor et al., 2021) and ADNI (Dadar et al., 2019) to assess WMHs in MCI and AD.

### Data availability statement

The data used for this analysis are available on request from the ADNI database (ida.loni.usc.edu).

### Statistical Analysis

Analyses were performed using MATLAB R2019b. Participant demographic information is presented in Table 1. Linear regression models were conducted to examine whether WMH, amyloid, and pTau would influence cognitive scores. Correction of multiple comparisons was completed using false discovery rate (FDR), p-values are reported as raw values with significance determined by FDR correction. *CognitiveScore_bl* represents executive function (or memory composite) score at baseline. For baseline measurements, 230 participants were included for both the executive functioning and memory composite models. Both CSF Aß42 and CSF pTau levels were downloaded from ADNI.

**Table 1:**
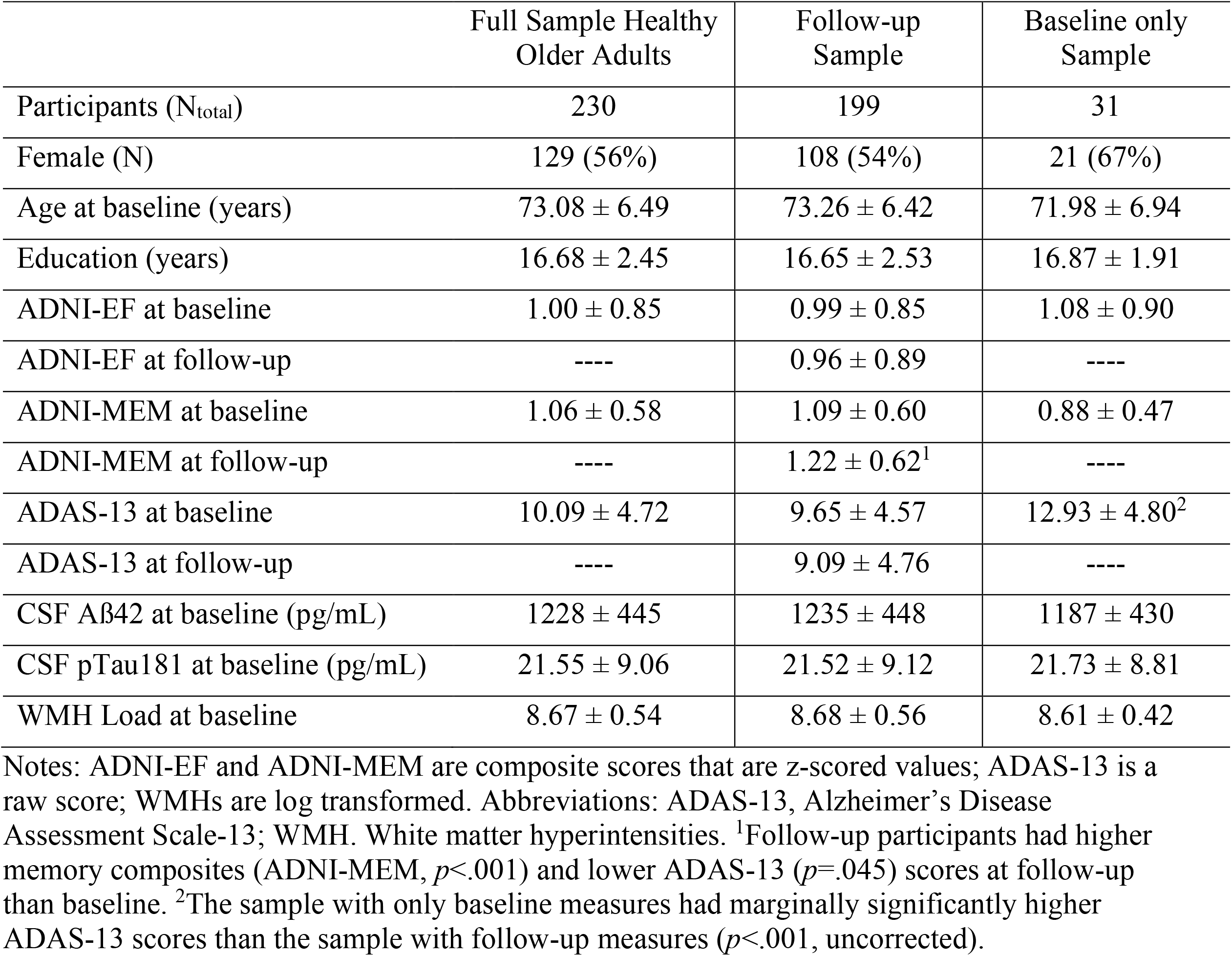
Descriptive statistics for the participants included in this study. Data are number (N) or mean ± standard deviation.

|  | Full Sample Healthy Older Adults | Follow-up Sample | Baseline only Sample |
| --- | --- | --- | --- |
| Participants (N<sub>total</sub>) | 230 | 199 | 31 |
| Female (N) | 129 (56%) | 108 (54%) | 21 (67%) |
| Age at baseline (years) | 73.08 ± 6.49 | 73.26 ± 6.42 | 71.98 ± 6.94 |
| Education (years) | 16.68 ± 2.45 | 16.65 ± 2.53 | 16.87 ± 1.91 |
| ADNI-EF at baseline | 1.00 ± 0.85 | 0.99 ± 0.85 | 1.08 ± 0.90 |
| ADNI-EF at follow-up | ---- | 0.96 ± 0.89 | ---- |
| ADNI-MEM at baseline | 1.06 ± 0.58 | 1.09 ± 0.60 | 0.88 ± 0.47 |
| ADNI-MEM at follow-up | ---- | 1.22 ± 0.62<sup>1</sup> | ---- |
| ADAS-13 at baseline | 10.09 ± 4.72 | 9.65 ± 4.57 | 12.93 ± 4.80<sup>2</sup> |
| ADAS-13 at follow-up | ---- | 9.09 ± 4.76 | ---- |
| CSF Aβ42 at baseline (pg/mL) | 1228 ± 445 | 1235 ± 448 | 1187 ± 430 |
| CSF pTau181 at baseline (pg/mL) | 21.55 ± 9.06 | 21.52 ± 9.12 | 21.73 ± 8.81 |
| WMH Load at baseline | 8.67 ± 0.54 | 8.68 ± 0.56 | 8.61 ± 0.42 |

Notes: ADNI-EF and ADNI-MEM are composite scores that are z-scored values; ADAS-13 is a raw score; WMHs are log transformed. Abbreviations: ADAS-13, Alzheimer's Disease Assessment Scale-13; WMH. White matter hyperintensities. <sup>1</sup>Follow-up participants had higher memory composites (ADNI-MEM, $p < .001$ ) and lower ADAS-13 ( $p = .045$ ) scores at follow-up than baseline. <sup>2</sup>The sample with only baseline measures had marginally significantly higher ADAS-13 scores than the sample with follow-up measures ( $p < .001$ , uncorrected).

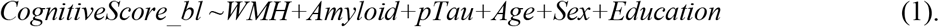

For follow-up measurements, 199 participants had follow-up scores over periods, 1-2 years (n=45), and 2-3 years (n=154) that were included for both the executive functioning and memory composite models. Baseline cognitive scores were also included in the follow-up model to ensure that the follow-up results account for more variance than what can be explained by the baseline scores.

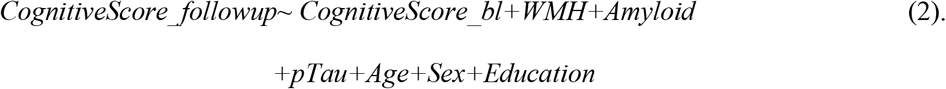

A linear regression was completed to examine whether amyloid and pTau influence white matter hyperintensities. All 230 participants were included in this analysis:

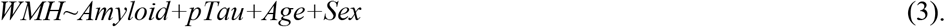

A linear regression was also completed to examine whether pTau and amyloid are associated. All 230 participants were included in this analysis:

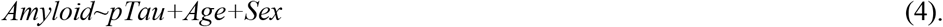

Additional t-tests were completed to compare the baseline characteristics of the 31 participants who did not have follow-up data to that of the 199 participants that had follow-up data to observe if there was evidence of selective attrition. The participants’ age, sex, education, CSF Aß42, CSF p-Tau181, ADNI-EF, ADNI-MEM, ADAS-13, and WMH load were compared between the two groups. At baseline, those that did not have follow-up scores had significantly increased ADAS-13 scores (*p*<.001, uncorrected); that is those who did not have follow-ups had lower performance at baseline in general cognitive functioning. No other group differences were significant (See Table 1).

## Results

### Cognitive Score Analysis

Figures 2 and 3 display executive functioning composite, memory composite, and ADAS-13 scores with Aß42, pTau, and WMH at baseline and follow-up. Table 2 summarizes the regression model results. At baseline, increased executive functioning scores were associated with higher CSF Aß42 (*t*= 3.58, *p<*.001) and higher education (*t*= 2.95, *p*=.003), and inversely associated with age (*t*= -5.63, *p<*.001). At follow-up, lower executive functioning scores were associated with increased WMHs (*t*= -2.84, *p*=.005), male sex (*t*= -2.22, *p*=.027), and increased age (*t*=-1.99, *p*=.049), however, age and sex were no longer significant after FDR correction. At follow-up, increased executive functioning scores were associated with increased education (*t*= 4.39, *p*<.001) and increased baseline executive functioning (*t*= 12.38, *p*<.001).

**Table 2:**
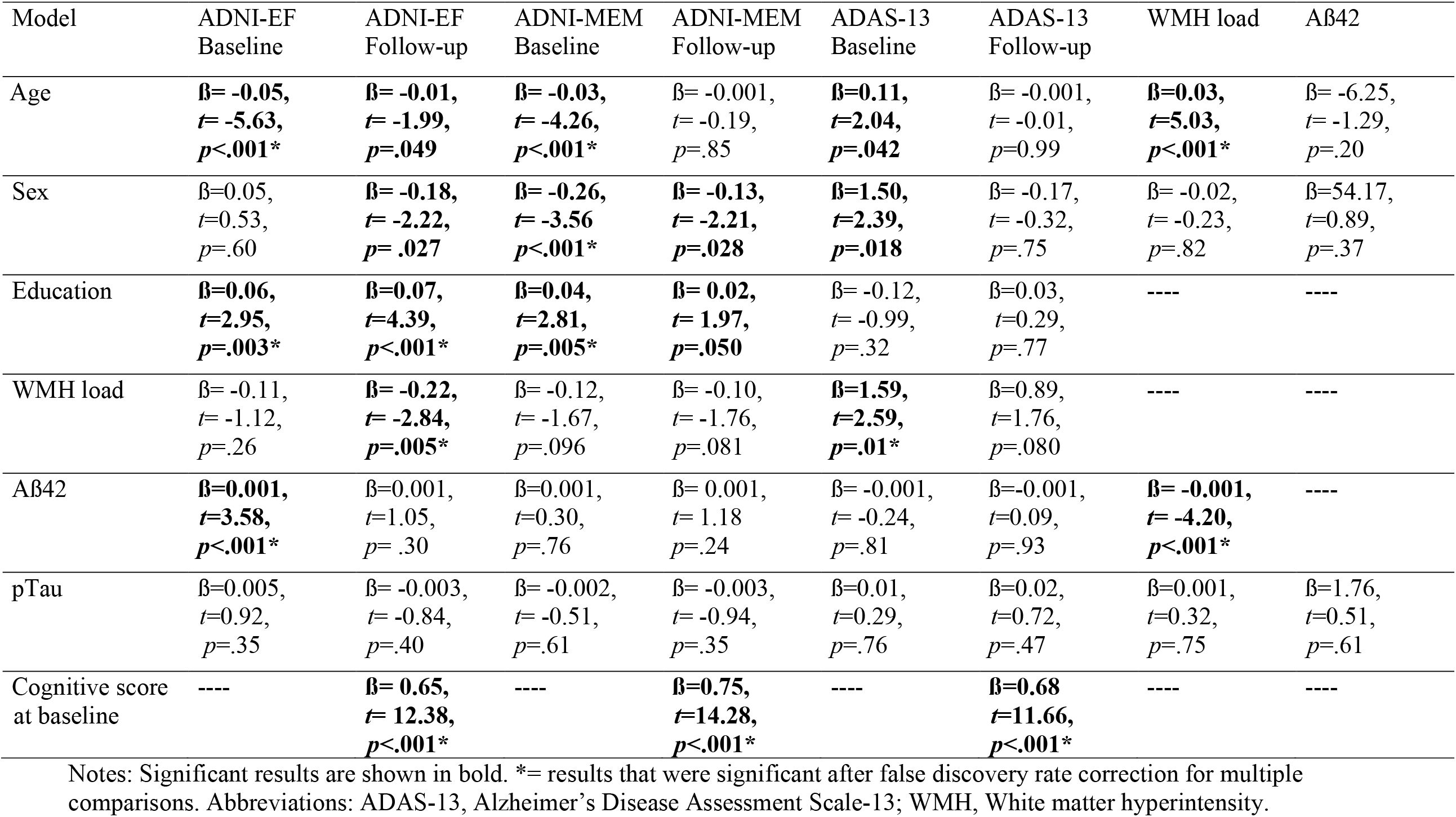
Regression model outputs.

| Model | ADNI-EF<br>Baseline | ADNI-EF<br>Follow-up | ADNI-MEM<br>Baseline | ADNI-MEM<br>Follow-up | ADAS-13<br>Baseline | ADAS-13<br>Follow-up | WMH load | Aβ42 |
| --- | --- | --- | --- | --- | --- | --- | --- | --- |
| Age | **β= -0.05,**<br>**t= -5.63,**<br>**p<.001\*** | **β= -0.01,**<br>**t= -1.99,**<br>**p=.049** | **β= -0.03,**<br>**t= -4.26,**<br>**p<.001\*** | β= -0.001,<br>t= -0.19,<br>p=.85 | **β=0.11,**<br>**t=2.04,**<br>**p=.042** | β= -0.001,<br>t= -0.01,<br>p=0.99 | **β=0.03,**<br>**t=5.03,**<br>**p<.001\*** | β= -6.25,<br>t= -1.29,<br>p=.20 |
| Sex | β=0.05,<br>t=0.53,<br>p=.60 | **β= -0.18,**<br>**t= -2.22,**<br>**p= .027** | **β= -0.26,**<br>**t= -3.56**<br>**p<.001\*** | **β= -0.13,**<br>**t= -2.21,**<br>**p=.028** | **β=1.50,**<br>**t=2.39,**<br>**p=.018** | β= -0.17,<br>t= -0.32,<br>p=.75 | β= -0.02,<br>t= -0.23,<br>p=.82 | β=54.17,<br>t=0.89,<br>p=.37 |
| Education | **β=0.06,**<br>**t=2.95,**<br>**p=.003\*** | **β=0.07,**<br>**t=4.39,**<br>**p<.001\*** | **β=0.04,**<br>**t=2.81,**<br>**p=.005\*** | **β= 0.02,**<br>**t= 1.97,**<br>**p=.050** | β= -0.12,<br>t= -0.99,<br>p=.32 | β=0.03,<br>t=0.29,<br>p=.77 | ---- | ---- |
| WMH load | β= -0.11,<br>t= -1.12,<br>p=.26 | **β= -0.22,**<br>**t= -2.84,**<br>**p=.005\*** | β= -0.12,<br>t= -1.67,<br>p=.096 | β= -0.10,<br>t= -1.76,<br>p=.081 | **β=1.59,**<br>**t=2.59,**<br>**p=.01\*** | β=0.89,<br>t=1.76,<br>p=.080 | ---- | ---- |
| Aβ42 | **β=0.001,**<br>**t=3.58,**<br>**p<.001\*** | β=0.001,<br>t=1.05,<br>p= .30 | β=0.001,<br>t=0.30,<br>p=.76 | β= 0.001,<br>t= 1.18<br>p=.24 | β= -0.001,<br>t= -0.24,<br>p=.81 | β=-0.001,<br>t=0.09,<br>p=.93 | **β= -0.001,**<br>**t= -4.20,**<br>**p<.001\*** | ---- |
| pTau | β=0.005,<br>t=0.92,<br>p=.35 | β= -0.003,<br>t= -0.84,<br>p=.40 | β= -0.002,<br>t= -0.51,<br>p=.61 | β= -0.003,<br>t= -0.94,<br>p=.35 | β=0.01,<br>t=0.29,<br>p=.76 | β=0.02,<br>t=0.72,<br>p=.47 | β=0.001,<br>t=0.32,<br>p=.75 | β=1.76,<br>t=0.51,<br>p=.61 |
| Cognitive score<br>at baseline | ---- | **β= 0.65,**<br>**t= 12.38,**<br>**p<.001\*** | ---- | **β=0.75,**<br>**t=14.28,**<br>**p<.001\*** | ---- | **β=0.68**<br>**t=11.66,**<br>**p<.001\*** | ---- | ---- |

Notes: Significant results are shown in bold. \*= results that were significant after false discovery rate correction for multiple comparisons. Abbreviations: ADAS-13, Alzheimer's Disease Assessment Scale-13; WMH, White matter hyperintensity.

**Figure 2:**
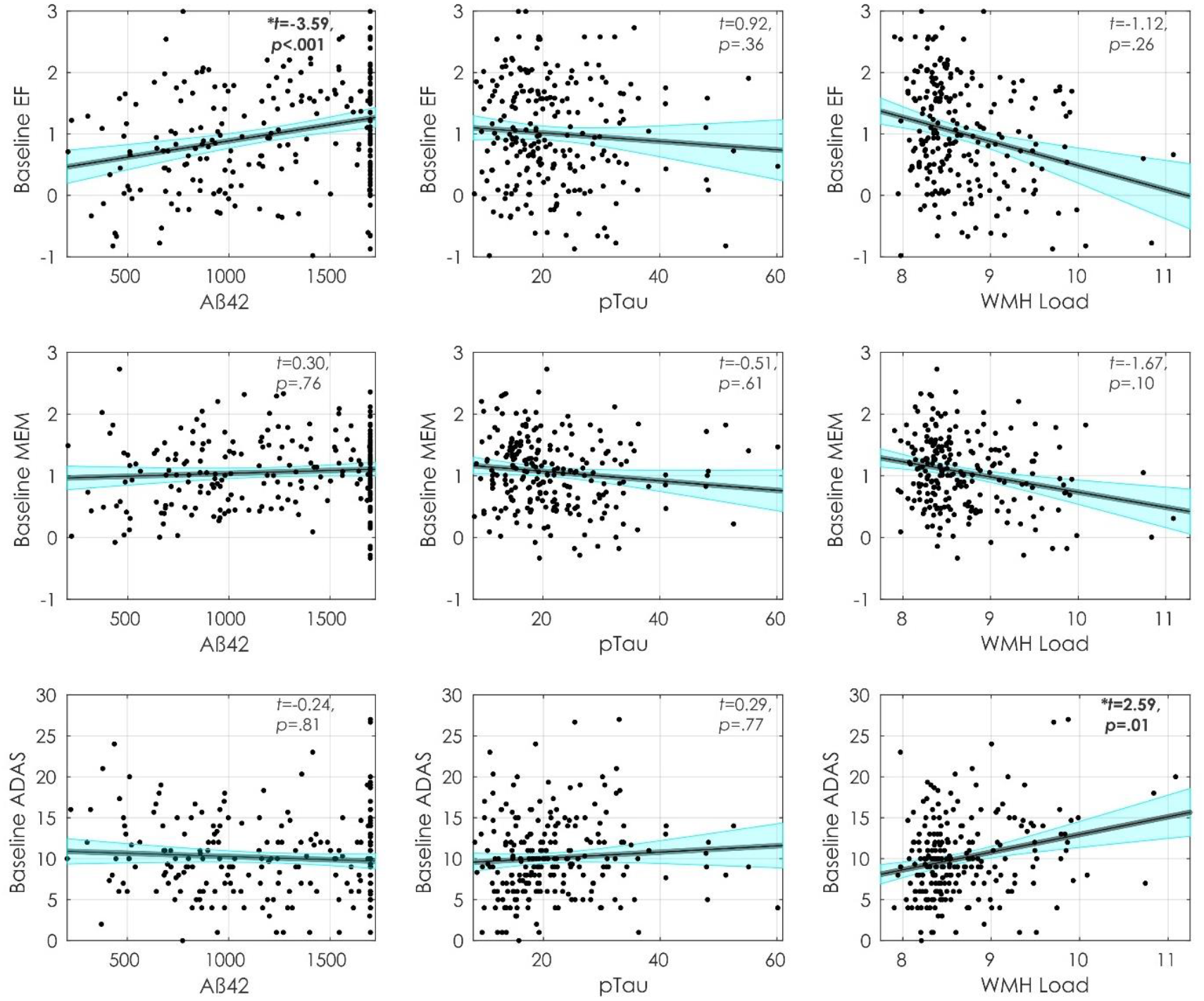
Baseline cognitive scores and their relationship with Aß42, pTau and WMH load. Figures show baseline executive functioning (first row), memory (second row) and ADAS-13 (third row) scores and their association with Aß42 (first column), pTau (Second column), and WMH Load (third column). Abbreviations: EF, executive functioning; MEM, memory; ADAS, Alzheimer’s Disease Assessment Scale; WMH, white matter hyperintensity.

**Figure 3:**
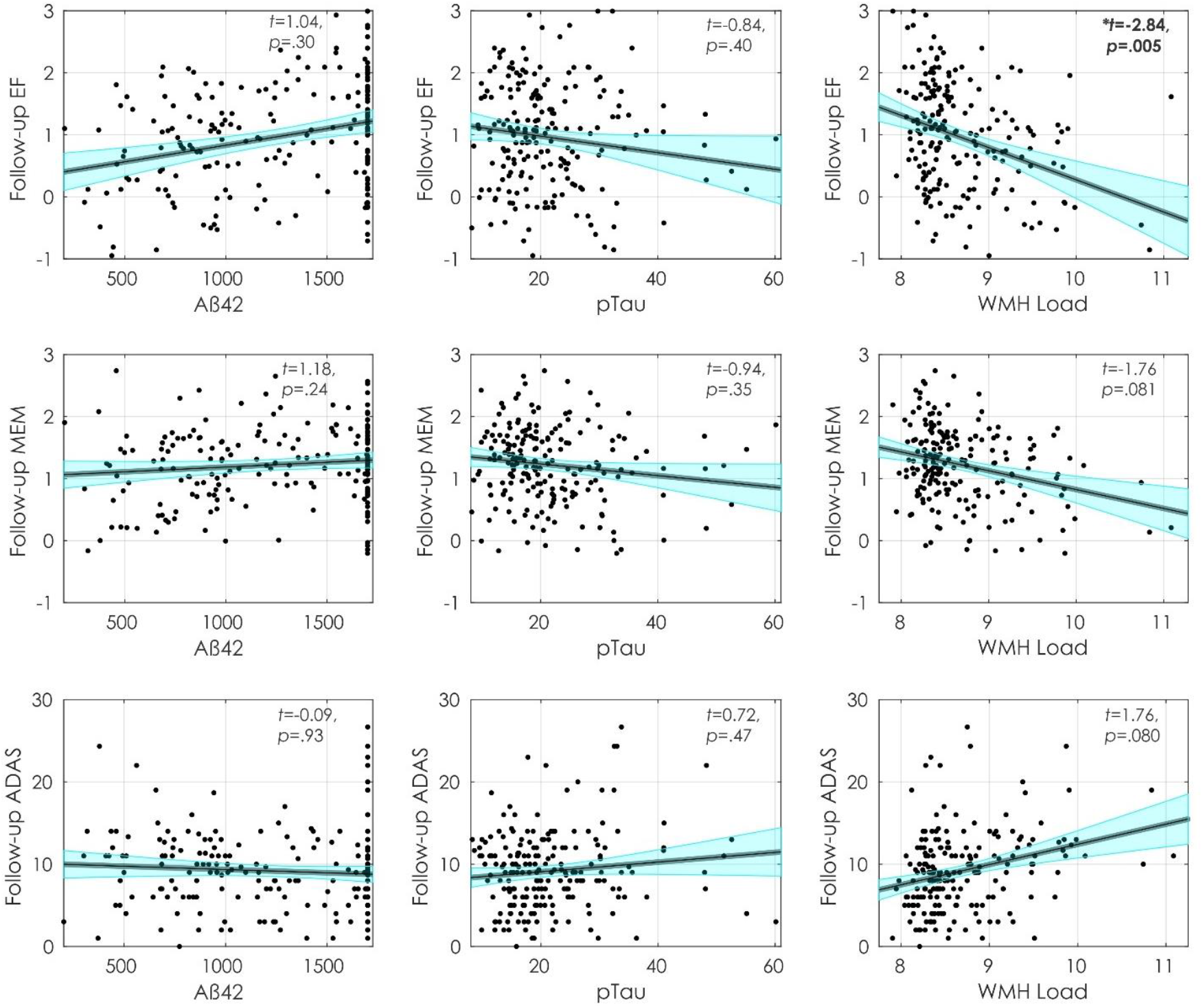
Follow-up cognitive scores and their relationship with Aß42, pTau and WMH load. Figures show follow-up executive functioning (first row), memory (second row) and ADAS-13 (third row) scores and their association with Aß42 (first column), pTau (Second column), and WMH Load (third column). Abbreviations: EF, executive functioning; MEM, memory; ADAS, Alzheimer’s Disease Assessment Scale; WMH, white matter hyperintensity.

At baseline, increased memory was associated with higher education (*t*= 2.81, *p*=.005). Lower memory scores were associated with increased age (*t*= -4.26, *p*<.001) and male sex (*t*= - 3.56, *p*<.001). At follow-up, increased memory was associated higher baseline memory (*t*= 14.28, *p*<.001). At follow-up increased memory was also associated with increased education (*t*=1.97, *p*=.05) and lower memory scores were associated with male sex (*t*= -2.21, *p*=.028) and marginally associated with increased WMHs (*t*=1.76, *p*=.081). However, the association between education, sex, and WMH with follow-up did not remain significant after FDR correction.

At baseline, higher ADAS-13 scores (i.e., worse performance) were associated with increased WMHs (*t*= 2.59, *p*=.01), age (*t*= 2.04, *p*=.042), and male sex (*t*= 2.39, *p*=.018); age and sex did were no longer significant after FDR correction. At follow-up, higher ADAS-13 scores were associated with higher baseline ADAS-13 scores (*t*= 11.66, *p*<.001) and marginally associated with increased WMHs (*t*= 1.76, *p*=.08).

### WMH, Tau, and Amyloid Analysis

Figure 4 displays the baseline associations between WMHs, CSF Aß42, and pTau. At baseline, WMHs were associated with increased CSF Aß42 (*t*= -4.20, *p*<.001) and age (*t*= 5.03, *p*<.001) but not pTau (*t*=0.32, *p*=0.75). CSF Aß42 was not observed to be associated with pTau (*t*= 0.51, *p*=.61), age (*t*= -1.29, *p*=.020) or male sex (*t*=0.89, *p*=0.37).

**Figure 4:**
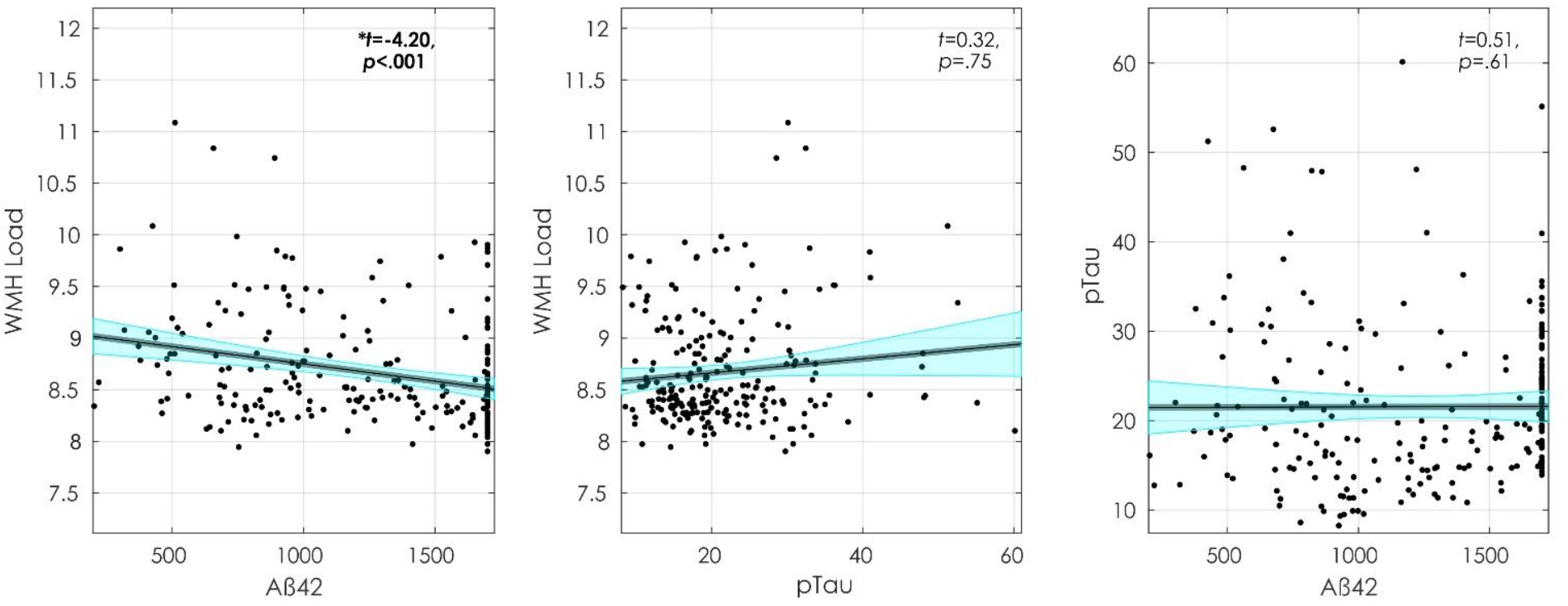
Baseline measurements of WMH load, Aß42, and pTau. Figures show, from left to right, WMH association with Aß42, WMH association with pTau, and pTau association with Aß42. Abbreviations: WMH, white matter hyperintensity.

## Discussion

The current study investigated the effects of WMH, CSF Aß42, and CSF pTau on current and future cognitive decline in cognitively normal older adults. We observed that CSF pTau was not associated with current or future decline for either memory or executive function in this cohort. CSF Aß42 burden was associated with only decreased baseline executive functioning. Increased WMH loads were associated with reduced performance on baseline global cognition (as measured by ADAS-13) and follow-up executive functioning. WMH loads were also related to baseline CSF Aß42 but not pTau. CSF Aß42 was not associated with CSF pTau. These results suggest WMH may contribute to the initial stages of cognitive decline in healthy older adults. While additional data and analysis are necessary, healthy older adults with WMH pathology may be less resilient to future pathology and at greater risk of cognitive impairment due to dementia than those without WMHs.

In this sample of cognitively unimpaired participants with no subjective cognitive decline, WMH loads appeared to have the largest association with cognitive changes in cognitively normal older adults. CSVD, as measured by WMHs, may be one of the first pathologies observed in the healthy older adult population that affects cognitive functioning. Pathologies that occur later in the aging process (i.e., tau and amyloid accumulation) may require less deposition to result in deterioration in cognitive functioning in those with high WMH loads. Reducing WMH may thus reduce the risk of future cognitive decline and the impact of other pathologies. The observation that future cognitive decline is impacted in healthy aging by WMH supports previous findings suggesting that WMHs increase the risk of MCI and dementia (Prins and Scheltens, 2015) by lowering the threshold for cognitive decline due to Alzheimer’s dementia. Risk factors that contribute to WMHs (e.g., hypertension and cardiovascular disease) are also associated with cognitive decline in healthy older adults (Marchant et al., 2012; Walker et al., 2017) and a higher conversion rate from MCI to AD (Ettorre et al., 2012). Taken together, these findings provide support an aggressive treatment strategy of vascular factors to help reduce future cognitive decline. For example, previous research has observed that people who undergo antihypertensive treatment exhibit less cognitive decline (Streit et al., 2019) and a reduced risk of dementia (Ding et al., 2020) than those with untreated high blood pressure. However, a recent meta-analysis suggests there is limited evidence to support the use of aggressive antihypertensive treatment to reduce cognitive decline (Dallaire-Théroux et al., 2021).

In MCI and AD, both PET and CSF tau and amyloid deposition is reported to be a strong predictor of cognitive decline (Bejanin et al., 2017; Lim et al., 2014; Rolstad et al., 2011). It is well-known that this accumulation of pathological amyloid and tau begins years before deficits in cognitive functioning can be detected (Leal et al., 2018), and thus may occur during healthy aging. The relationship between tau and amyloid in healthy older adults is, however, less prominent. Nevertheless, some associations between CSF tau and amyloid (Donohue et al., 2017; Stomrud et al., 2010; Verberk et al., 2020) with reduced cognitive performance in healthy older adults have been observed. CSF pTau was not associated with cognitive decline in our sample of healthy older adults without subjective cognitive decline. The sample of participants in this study comprises healthy older adults who exhibit low overall pTau burdens making it difficult to detect associations between tau and cognition at this stage (Jack et al., 2017). Amyloid deposition may occur on average 13.3 years earlier than tau accumulation (Therneau et al., 2021). Thus, it is not surprising that Aß42 was associated with baseline executive functioning in the absence of associations between pTau and cognition. The fastest rates of cognitive decline are observed in those who have both elevated amyloid and tau deposition (Sperling et al., 2019). Therefore, the low pTau levels in our sample may have influenced why we did not observe an association between pTau and cognition at baseline or follow-up or between Aß42 and cognition at follow-up in our healthy older adult sample.

The participants who did not have follow-up data had higher ADAS-13 scores (i.e., more cognitive impairment) at baseline than those who had follow-up data. This finding suggests that those participants may be closer to the dementia/AD trajectory. In the current study, we only have 2 to 3 years of follow-up data so we cannot examine differences in people who develop more advanced accumulation or convert to dementia/AD. Future research should examine longer follow-ups to determine whether cognitively normal healthy older adults with WMHs have a faster rate of cognitive decline, if they are more likely to convert to dementia/AD, and if elevated amyloid and tau contribute to this decline. In this study, while we did observe a relationship between WMH and amyloid at baseline we did not observe a relationship between WMH and tau. This finding is similar to a previous study that reported no associations between tau-PET and WMH burden, but reported that amyloid-PET load correlated with WMH (Graff-Radford et al., 2019).

The image processing methods employed in this study have been developed and extensively validated for use in multi-center and multi-scanner studies. These processing methods have previously observed WMH effects in PD (Dadar et al., 2021, 2020a), healthy aging and AD (Dadar et al., 2020a). Therefore, the lack of association between WMH with amyloid, tau, and some cognitive scores is not the result of poor sensitivity of the image processing methods employed.

The limitations of the current study include the relatively short follow-up period, limited sample size, and high participant education. Our follow-up times were limited to 3 years because we included participants from ADNI-3. Longer follow-ups may demonstrate stronger associations between tau and amyloid pathologies and cognitive decline. The relatively limited sample in this study did not allow for the testing of the interaction between amyloid, tau, and WMH on cognition with enough statistical power. The high education of this sample may limit the generalizability of these results to other populations. Some research has shown that WMHs may also be associated with other pathologies than cerebrovascular disease (McAleese et al., 2021). However, the methodologies needed to differentiate these pathologies are unavailable in this dataset, limiting our ability to identify the cause of the WMHs. Future research is needed to examine the underlying pathologies associated with WMHs.

## Conclusion

The current study sought to examine the relationship between CSVD as measured by WMH load, with amyloid, and tau measured with CSF, and their effects on cognitive functioning in healthy older adults without subjective cognitive decline. Our findings suggest that in healthy older adults who are not likely to be in the preclinical AD stage, as reflected by low tau levels, WMHs may be one of the first pathologies to contribute to age-related cognitive decline. It is possible that WMH burden may preceded tau and amyloid deposition, however, without being able to determine who converts to AD, this hypothesis needs to be further examined. Despite being unable to determine which participants will eventually develop dementia in this study, our findings that WMHs contribute to cognitive decline aligns with previous work suggesting that vascular biomarkers may improve our current understanding of AD pathophysiology and dementia (Sweeney et al., 2019).

## Acknowledgments

Data collection and sharing for this project was funded by the Alzheimer’s Disease Neuroimaging Initiative (ADNI) (National Institutes of Health Grant U01 AG024904) and DOD ADNI (Department of Defense award number W81XWH-12-2-0012). ADNI is funded by the National Institute on Aging, the National Institute of Biomedical Imaging and Bioengineering, and through generous contributions from the following: AbbVie, Alzheimer’s Association; Alzheimer’s Drug Discovery Foundation; Araclon Biotech; BioClinica, Inc.; Biogen; Bristol-Myers Squibb Company; CereSpir, Inc.; Cogstate; Eisai Inc.; Elan Pharmaceuticals, Inc.; Eli Lilly and Company; EuroImmun; F. Hoffmann-La Roche Ltd and its affiliated company Genentech, Inc.; Fujirebio; GE Healthcare; IXICO Ltd.; Janssen Alzheimer Immunotherapy Research & Development, LLC.; Johnson & Johnson Pharmaceutical Research & Development LLC.; Lumosity; Lundbeck; Merck & Co., Inc.; Meso Scale Diagnostics, LLC.; NeuroRx Research; Neurotrack Technologies; Novartis Pharmaceuticals Corporation; Pfizer Inc.; Piramal Imaging; Servier; Takeda Pharmaceutical Company; and Transition Therapeutics. The Canadian Institutes of Health Research is providing funds to support ADNI clinical sites in Canada. Private sector contributions are facilitated by the Foundation for the National Institutes of Health (www.fnih.org). The grantee organization is the Northern California Institute for Research and Education, and the study is coordinated by the Alzheimer’s Therapeutic Research Institute at the University of Southern California. ADNI data are disseminated by the Laboratory for Neuro Imaging at the University of Southern California.

## Notes

**Funding information** Alzheimer’s Disease Neuroimaging Initiative; This research was supported by a grant from the Canadian Institutes of Health Research and The Louise & André Charron Family.

**Financial Disclosures** Dr. Morrison is supported by a Canadian Institute of Health Research Postdoctoral fellowship. Dr. Dadar is supported by a scholarship from the Canadian Consortium on Neurodegeneration in Aging as well as an Alzheimer Society Research Program (ASRP) postdoctoral award. The Consortium is supported by a grant from the Canadian 24 Institutes of Health Research with funding from several partners including the Alzheimer Society of Canada, Sanofi, and Women’s Brain Health Initiative.

Dr. Villeneuve reports work supported by a Canada Research Chair, a Canadian Institutes of Health Research Foundation Grant, a Canada Fund for Innovation Grant, an Alzheimer’s Association Grant, and an Alzheimer’s society of Canada and Fonds de recherche Sante Quebec fellowship.

### Competing Interest Statement

The authors have declared no competing interest.

## References

Anor CJ, Dadar M, Collins DL, Tartaglia MC., 2021. The Longitudinal Assessment of Neuropsychiatric Symptoms in Mild Cognitive Impairment and Alzheimer’s Disease and Their Association With White Matter Hyperintensities in the National Alzheimer’s Coordinating Center’s Uniform Data Set. Biol Psychiatry Cogn Neurosci Neuroimaging 6(1), 70–78. doi:10.1016/j.bpsc.2020.03.006

Aschenbrenner AJ, Gordon BA, Benzinger TLS, Morris JC, Hassenstab JJ., 2018. Influence of tau PET, amyloid PET, and hippocampal volume on cognition in Alzheimer disease. Neurology 91(9), e859–e866. doi:10.1212/WNL.0000000000006075

Avants BB, Epstein CL, Grossman M, Gee JC., 2008. Symmetric diffeomorphic image registration with cross-correlation: Evaluating automated labeling of elderly and neurodegenerative brain. Med Image Anal 12(1), 26–41. doi:10.1016/j.media.2007.06.004

Bejanin A, Schonhaut, D.R., La Joie, R., Kramer, J.H., Baker, S.L., Sosa, N., Ayakta, N., Cantwell, A., Janabi, M., Lauriola, M., O’Neil, J.P., Gorno-Tempini, M.L., Miller, Z.A., Rosen, H.J., Miller, B.L., Jagust, W.J., Rabinovici, G.D., 2017. Tau pathology and neurodegeneration contribute to cognitive impairment in Alzheimer’s disease. Brain 140(12), 3286–3300. doi:https://dx.doi.org/10.1093/brain/awx243

Boyle PA, Yu, L., Fleischman, D.A., Leurgans, S., Yang, J., Wilson, R.S., Schneider, J.A., Arvanitakis, Z., Arfanakis, K., Bennett, D.A., 2016. White matter hyperintensities, incident mild cognitive impairment, and cognitive decline in old age. Ann Clin Transl Neurol 3(10), 791–800. doi:10.1002/acn3.343

Coupe P, Yger P, Prima S, Hellier P, Kervrann C, Barillot C., 2008 An optimized blockwise nonlocal means denoising filter for 3-D magnetic resonance images. IEEE Trans Med Imaging 27(4), 425–441. doi:10.1109/TMI.2007.906087

Craig-schapiro R, Fagan AM, Holtzman DM., 2009 Biomarkers of Alzheimer’s disease. Neurobiol Dis 35(2), 128–140. doi:10.1016/j.nbd.2008.10.003

Crane PK, Carle, A., Gibbons, L.E., Insel, P., Mackin, R.S., Gross, A., Jones, R.N., Mukherjee, S., Curtis, S.M., Harvey, D., Weiner, M., 2012. Development and assessment of a composite score for memory in the Alzheimer’s Disease Neuroimaging Initiative (ADNI). Brain Imaging Behav 6(4), 502–516. doi:10.1007/s11682-012-9186-z

Dadar, M., Camicioli, R., Duchesne, S., Collins, D.L., 2020a. The temporal relationships between white matter hyperintensities, neurodegeneration, amyloid beta, and cognition. Alzheimer’s Dement. Diagnosis, Assess. Dis. Monit. 1, e12091. https://doi.org/10.1002/dad2.12091

Dadar, M., Fereshtehnejad, S.M., Zeighami, Y., Dagher, A., Postuma, R.B., Collins, D.L., 2020b. White Matter Hyperintensities Mediate Impact of Dysautonomia on Cognition in Parkinson’s Disease. Mov. Disord. Clin. Pract. 7(6), 639–647. https://doi.org/10.1002/mdc3.13003

Dadar, M., Fonov, V.S., Collins, D.L., 2018a. A comparison of publicly available linear MRI stereotaxic registration techniques. Neuroimage 174, 191–200. https://doi.org/10.1016/j.neuroimage.2018.03.025

Dadar, M., Gee, M., Shuaib, A., Duchesne, S., Camicioli, R., 2020c. Cognitive and motor correlates of grey and white matter pathology in Parkinson’s disease. NeuroImage Clin. 27, 102353. https://doi.org/10.1016/j.nicl.2020.102353

Dadar, M., Maranzano, J., Ducharme, S., Collins, D.L., 2019. White matter in different regions evolves differently during progression to dementia. Neurobiol. Aging 76, 71–79. https://doi.org/10.1016/j.neurobiolaging.2018.12.004

Dadar, M., Misquitta, K., Anor, C.J., Fonov, V.S., Tartaglia, M.C., Carmichael, O.T., Decarli, C., Collins, D.L., 2017a. NeuroImage Performance comparison of 10 different classification techniques in segmenting white matter hyperintensities in aging 157, 233–249. https://doi.org/10.1016/j.neuroimage.2017.06.009

Dadar, M., Miyasaki, J., Duchesne, S., Camicioli, R., 2021. White matter hyperintensities mediate the impact of amyloid ß on future freezing of gait in Parkinson’s disease. Park. Relat. Disord. 85, 95–101. https://doi.org/10.1016/j.parkreldis.2021.02.031

Dadar, M., Pascoal, T.A., Manitsirikul, S., Misquitta, K., Fonov, V.S., Carmela, M., Breitner, J., Rosa-neto, P., Carmichael, O.T., Decarli, C., Collins, D.L., 2017b. Validation of a Regression Technique for Segmentation of White Matter Hyperintensities in Alzheimer’s Disease 36(8), 1758–1768.

Dadar, M., Zeighami, Y., Yau, Y., Fereshtehnejad, S.M., Maranzano, J., Postuma, R.B., Dagher, A., Collins, D.L., 2018b. White matter hyperintensities are linked to future cognitive decline in de novo Parkinson’s disease patients. NeuroImage Clin. 20, 892–900. https://doi.org/10.1016/j.nicl.2018.09.025

Dallaire-Théroux, C., Quesnel-Olivo, M. H., Brochu, K., Bergeron, F., O’Connor, S., Turgeon, A. F., … & Duchesne, S. (2021). Evaluation of Intensive vs Standard Blood Pressure Reduction and Association With Cognitive Decline and Dementia: A Systematic Review and Meta-analysis. JAMA Network Open, 4(11), e2134553–e2134553. doi:10.1001/jamanetworkopen.2021.34553

Ding, J., Davis-Plourde, K.L., Sedaghat, S., Tully, P.J., Wang, W., Phillips, C., Pase, M.P., Himali, J.J., Gwen Windham, B., Griswold, M., Gottesman, R., Mosley, T.H., White, L., Guðnason, V., Debette, S., Beiser, A.S., Seshadri, S., Ikram, M.A., Meirelles, O., Tzourio, C., Launer, L.J., 2020. Antihypertensive medications and risk for incident dementia and Alzheimer’s disease: a meta-analysis of individual participant data from prospective cohort studies. Lancet Neurol. 19, 61–70. https://doi.org/10.1016/S1474-4422(19)30393-X

Donohue, M.C., Sperling, R.A., Petersen, R., Sun, C.K., Weiner, M., Aisen, P.S., 2017. Association between elevated brain amyloid and subsequent cognitive decline among cognitively normal persons. JAMA - J. Am. Med. Assoc. 317, 2305–2316. https://doi.org/10.1001/jama.2017.6669

Ettorre, E., Cerra, E., Marigliano, B., Vigliotta, M., Vulcano, A., Fossati, C., De Benedetto, G., Servello, A., Andreozzi, P., Marigliano G., Servello, A.; Andreozzi, P., 2012. Role of cardiovascular risk factors (CRF) in the patients with mild cognitive impairment (MCI). Arch. Gerontol. Geriatr. 54, 330–332. https://doi.org/http://dx.doi.org/10.1016/j.archger.2011.04.025

Gibbons, L.E., Carle, A.C., Mackin, R.S., Harvey, D., Mukherjee, S., Insel, P., Curtis, S.M., Mungas, D., Crane, P.K., 2012. A composite score for executive functioning, validated in Alzheimer’s Disease Neuroimaging Initiative (ADNI) participants with baseline mild cognitive impairment. Brain Imaging Behav. 6, 517–527. https://doi.org/10.1007/s11682-012-9176-1

Graff-Radford, J., Arenaza-Urquijo, E.M., Knopman, D.S., Schwarz, C.G., Brown, R.D., Rabinstein, A.A., Gunter, J.L., Senjem, M.L., Przybelski, S.A., Lesnick, T., Ward, C., Mielke, M.M., Lowe, V.J., Petersen, R.C., Kremers, W.K., Kantarci, K., Jack, C.R., Vemuri, P., 2019. White matter hyperintensities: Relationship to amyloid and tau burden. Brain 142(8), 2483–2491. https://doi.org/10.1093/brain/awz162

Hedden, T., Mormino, E.C., Amariglio, R.E., Younger, A.P., Schultz, A.P., Becker, J.A., Buckner, R.L., Johnson, K.A., Sperling, R.A., Rentz, D.M., 2012. Cognitive profile of amyloid burden and white matter hyperintensities in cognitively normal older adults. J. Neurosci. 32(46), 16233–16242. https://doi.org/10.1523/JNEUROSCI.2462-12.2012

Jack, C.R., Wiste, H.J., Weigand, S.D., Therneau, T.M., Lowe, V.J., Knopman, D.S., Gunter, J.L., Senjem, M.L., Jones, D.T., Kantarci, K., Machulda, M.M., Mielke, M.M., Roberts, R.O., Vemuri, P., Reyes, D.A., Petersen, R.C., 2017. Defining imaging biomarker cut points for brain aging and Alzheimer’s disease. Alzheimer’s Dement. 13(3), 205–216. https://doi.org/10.1016/j.jalz.2016.08.005

Kaskikallio, A., Karrasch, M., Koikkalainen, J., Lötjönen, J., Rinne, J.O., Tuokkola, T., Parkkola, R., Grönholm-Nyman, P., 2020. White Matter Hyperintensities and Cognitive Impairment in Healthy and Pathological Aging: A Quantified Brain MRI Study. Dement. Geriatr. Cogn. Disord. 48(5-6) 297–307. https://doi.org/10.1159/000506124

Kloppenborg, R.P., Geerlings, M.I., 2014. Presence and progression of white matter hyperintensities and cognition. Neurology 82(23),2127–2138. https://doi.org/10.1212/WNL.0000000000000505

Leal, S.L., Lockhart, S.N., Maass, A., Bell, R.K., Jagust, W.J., 2018. Subthreshold amyloid predicts tau deposition in aging. J. Neurosci. 38(19), 4482–4489. https://doi.org/10.1523/JNEUROSCI.0485-18.2018

Lim, Y.Y., Maruff, P., Pietrzak, R.H., Ames, D., Ellis, K.A., Harrington, K., Lautenschlager, N.T., Szoeke, C., Martins, R.N., Masters, C.L., Villemagne, V.L., Rowe, C.C., 2014. Effect of amyloid on memory and non-memory decline from preclinical to clinical Alzheimer’s disease. Brain 137(1), 221–231. https://doi.org/10.1093/brain/awt286

Malpas, C.B., Sharmin, S., Kalincik, T., 2021. The histopathological staging of tau, but not amyloid, corresponds to antemortem cognitive status, dementia stage, functional abilities and neuropsychiatric symptoms. Int. J. Neurosci. 131(8), 800–809. https://doi.org/10.1080/00207454.2020.1758087

Manera, A.L., Dadar, M., Fonov, V., Collins, D.L., 2020. CerebrA, registration and manual label correction of Mindboggle-101 atlas for MNI-ICBM152 template. Sci. Data 1–9. https://doi.org/10.1038/s41597-020-0557-9

Marchant, N.L., Reed, B.R., DeCarli, C.S., Madison, C.M., Weiner, M.W., Chui, H.C., Jagust, W.J., 2012. Cerebrovascular disease, beta-amyloid, and cognition in aging. Neurobiol. Aging 33(5), 1006.e25-1006.e36. https://doi.org/10.1016/j.neurobiolaging.2011.10.001

McAleese, K. E., Miah, M., Graham, S., Hadfield, G. M., Walker, L., Johnson, M., … & Attems, J. (2021). Frontal white matter lesions in Alzheimer’s disease are associated with both small vessel disease and AD-associated cortical pathology. Acta Neuropathologica, 142(6), 937–950. https://doi.org/10.1007/s00401-021-02376-2

Meyer, P.F., Pichet Binette, A., Gonneaud, J., Breitner, J.C.S., Villeneuve, S., 2020. Characterization of Alzheimer Disease Biomarker Discrepancies Using Cerebrospinal Fluid Phosphorylated Tau and AV1451 Positron Emission Tomography. JAMA Neurol. 77(4), 508–516. https://doi.org/10.1001/jamaneurol.2019.4749

Mohs, R.C., Knopman, D.S., Petersen, R.C., Ferris, S.H., Ernesto, C., Grundman, M., Sano, M., Bieliauskas, L., Geldmacher, D.S., Clark, C., Thal, L., 1997. Development of cognitive instruments for use in clinical trials of antidementia drugs: Additions to the Alzheimer’s Disease Assessment Scale that broaden its scope. Alzheimer Dis. Assoc. Disord. 11(Suppl2), S13–S21.

Olsson, A., Vanderstichele, H., Andreasen, N., De Meyer, G., Wallin, A., Holmberg, B., Rosengren, L., Vanmechelen, E., Blennow, K., 2005. Simultaneous measurement of β-amyloid(1-42), total Tau, and phosphorylated Tau (Thr181) in cerebrospinal fluid by the xMAP technology. Clin. Chem. 51(2), 336–345. https://doi.org/10.1373/clinchem.2004.039347

Perrin, R., Fagan, A.M., Holtzman, D.M., 2009. Multi-modal techniques for diagnosis and prognosis of Alzheimer’s disease. Nature 461(7266), 916–922. https://doi.org/10.1038/nature08538.Multi-modal

Prins, N.D., Scheltens, P., 2015. White matter hyperintensities, cognitive impairment and dementia: An update. Nat. Rev. Neurol. 11(3), 157–165. https://doi.org/10.1038/nrneurol.2015.10

Prins, N.D., Van Dijk, E.J., Den Heijer, T., Vermeer, S.E., Koudstaal, P.J., Oudkerk, M., Hofman, A., Breteler, M.M.B., 2004. Cerebral white matter lesions and the risk of dementia. Arch. Neurol. 61(10), 1531–1534. https://doi.org/10.1001/archneur.61.10.1531

Rabin, L.A., Smart, C.M., Amariglio, R.E., 2017. Subjective Cognitive Decline in Preclinical Alzheimer’s Disease. Annu. Rev. Clin. Psychol. 13, 369–396. https://doi.org/10.1146/annurev-clinpsy-032816-045136

Rhodius-Meester, H.F.M., Benedictus, M.R., Wattjes, M.P., Barkhof, F., Scheltens, P., Muller, M., van der Flier, W.M., 2017. MRI visual ratings of brain atrophy and white matter hyperintensities across the spectrum of cognitive decline are differently affected by age and diagnosis. Front. Aging Neurosci. 9, 1–12. https://doi.org/10.3389/fnagi.2017.00117

Rolstad, S., Berg, A.I., Bjerke, M., Blennow, K., Johansson, B., Zetterberg, H., Wallin, A., 2011. Amyloid-beta42 is associated with cognitive impairment in healthy elderly and subjective cognitive impairment. J. Alzheimer’s Dis. 26(1), 135–142.

Sanford, R., Strain, J., Dadar, M., Maranzano, J., Bonnet, A., Mayo, N.E., Scott, S.C., Fellows, L.K., Ances, B.M., Collins, D.L., Neurologie, D., Cedex, R., 2019. HIV infection and cerebral small vessel disease are independently associated with brain atrophy and cognitive impairment 33(7), 1197–1205. https://doi.org/10.1097/QAD.0000000000002193.HIV

Schindler, S.E., Gray, J.D., Gordon, B.A., Xiong, C., Batrla-Utermann, R., Quan, M., Wahl, S., Benzinger, T.L.S., Holtzman, D.M., Morris, J.C., Fagan, A.M., 2018. Cerebrospinal fluid biomarkers measured by Elecsys assays compared to amyloid imaging. Alzheimer’s Dement. 14(11), 1460–1469. https://doi.org/10.1016/j.jalz.2018.01.013

Schöll, M., Lockhart, S.N., Schonhaut, D.R., O’Neil, J.P., Janabi, M., Ossenkoppele, R., Baker, S.L., Vogel, J.W., Faria, J., Schwimmer, H.D., Rabinovici, G.D., Jagust, W.J., 2016. PET Imaging of Tau Deposition in the Aging Human Brain. Neuron 89(5), 971–982. https://doi.org/10.1016/j.neuron.2016.01.028

Shaw, L.M., Vanderstichele, H., Knapik-Czajka, M., Clark, C.M., Aisen, P.S., Petersen, R.C., Blennow, K., Soares, H., Simon, A., Lewczuk, P., Dean, R., Siemers, E., Potter, W., Lee, V.M.Y., Trojanowski, J.Q., 2009. Cerebrospinal fluid biomarker signature in alzheimer’s disease neuroimaging initiative subjects. Ann. Neurol. 65(4), 403–413. https://doi.org/10.1002/ana.21610

Sled, J.G., Zijdenbos, A.P., Evans, A.C., 1998. A nonparametric method for automatic correction of intensity nonuniformity in mri data. IEEE Trans. Med. Imaging 17(1), 87–97. https://doi.org/10.1109/42.668698

Sperling, R.A., Aisen, P.S., Beckett, L.A., Bennett, D.A., Craft, S., Fagan, A.M., Iwatsubo, T., Jack, C.R., Kaye, J., Montine, T.J., Park, D.C., Reiman, E.M., Rowe, C.C., Siemers, E., Stern, Y., Yaffe, K., Carrillo, M.C., Thies, B., Morrison-Bogorad, M., Wagster, M. V., Phelps, C.H., 2011. Toward defining the preclinical stages of Alzheimer’s disease: Recommendations from the National Institute on Aging-Alzheimer’s Association workgroups on diagnostic guidelines for Alzheimer’s disease. Alzheimer’s Dement. 7(3), 280–292. https://doi.org/10.1016/j.jalz.2011.03.003

Sperling, R.A., Mormino, E.C., Schultz, A.P., Betensky, R.A., Papp, K. V., Amariglio, R.E., Hanseeuw, B.J., Buckley, R., Chhatwal, J., Hedden, T., Marshall, G.A., Quiroz, Y.T., Donovan, N.J., Jackson, J., Gatchel, J.R., Rabin, J.S., Jacobs, H., Yang, H.S., Properzi, M., Kirn, D.R., Rentz, D.M., Johnson, K.A., 2019. The impact of amyloid-beta and tau on prospective cognitive decline in older individuals. Ann. Neurol. 85(2), 181–193. https://doi.org/10.1002/ana.25395

Stomrud, E., Hansson, O., Zetterberg, H., Blennow, K., Minthon, L., Londos, E., 2010. Correlation of longitudinal cerebrospinal fluid biomarkers with cognitive decline in healthy older adults. Arch. Neurol. 67(2), 217–223. https://doi.org/10.1001/archneurol.2009.316

Streit, S., Poortvliet, R.K.E., Den Elzen, W.P.J., Blom, J.W., Gussekloo, J., 2019. Systolic blood pressure and cognitive decline in older adults with hypertension. Ann. Fam. Med. 17(2), 100–107. https://doi.org/10.1370/afm.2367

Sutphen, C. L., McCue, L., Herries, E. M., Xiong, C., Ladenson, J. H., Holtzman, D. M., & Fagan, A. M. (2018). Longitudinal decreases in multiple cerebrospinal fluid biomarkers of neuronal injury in symptomatic late onset Alzheimer’s disease. Alzheimer’s & Dementia, 14(7), 869–879.

Sweeney, M.D., Montagne, A., Sagare, A.P., Nation, D.A., Schneider, L.S., Chui, H.C., Harrington, M.G., Pa, J., Law, M., Wang, D.J. and Jacobs, R.E., 2019. Vascular dysfunction—the disregarded partner of Alzheimer’s disease. Alzheimer’s & Dementia, 15(1), 158–167.

Therneau, T.M., Knopman, D.S., Lowe, V.J., Botha, H., Graff-Radford, J., Jones, D.T., Vemuri, P., Mielke, M.M., Schwarz, C.G., Senjem, M.L. and Gunter, J.L., 2021. Relationships between β-amyloid and tau in an elderly population: An accelerated failure time model. Neuroimage, 242, p.118440. doi:10.1016/j.neuroimage.2021.118440

Vemuri, P., Lesnick, T.G., Przybelski, S.A., Knopman, D.S., Preboske, G.M., Kantarci, K., Raman, M.R., Machulda, M.M., Mielke, M.M., Lowe, V.J., Senjem, M.L., Gunter, J.L., Rocca, W.A., Roberts, R.O., Petersen, R.C., Jack, C.R., 2015. Vascular and amyloid pathologies are independent predictors of cognitive decline in normal elderly. Brain 138(3), 761–771. https://doi.org/10.1093/brain/awu393

Verberk, I.M.W., Hendriksen, H.M.A., van Harten, A.C., Wesselman, L.M.P., Verfaillie, S.C.J., van den Bosch, K.A., Slot, R.E.R., Prins, N.D., Scheltens, P., Teunissen, C.E., Van der Flier, W.M., 2020. Plasma amyloid is associated with the rate of cognitive decline in cognitively normal elderly: the SCIENCe project. Neurobiol. Aging 89, 99–107. https://doi.org/10.1016/j.neurobiolaging.2020.01.007

Walker, K.A., Power, M.C., Gottesman, R.F., 2017. Defining the Relationship Between Hypertension, Cognitive Decline, and Dementia: a Review. Curr. Hypertens. Rep. 19(3), 24. https://doi.org/10.1007/s11906-017-0724-3

